# Phosphorylation of spleen tyrosine kinase Y130 positively regulates intracellular signaling and functional responses in platelets

**DOI:** 10.64898/2026.08.31.748427

**Authors:** Manal Elzoheiry, Carol Dangelmaier, Dhruv N Vajipayajula, Monica Wright, Peter Coopman, Alexander Y Tsygankov, Satya P Kunapuli

## Abstract

Syk is a non-receptor type protein-tyrosine kinase (PTK), which is associated with platelets surface receptors, glycoprotein VI (GPVI) and C-type lectin-like receptor II-type (CLEC-2). Syk is also expressed in most hematopoietic lineage cells and other cells, such as fibroblasts and neuronal cells. Syk has two tandem SH2 motifs and a C-terminal kinase domain, which are interrupted by interdomains A and B containing multiple tyrosine residues playing a regulatory role upon phosphorylation. This study aims to evaluate the role of Y130 in Syk signaling in platelets. Syk(Y130F) knock-in (KI) mice we generated using the CRISPR-Cas9 technique represent the first *in-vivo* model harboring this mutation. Using this system, we compared the platelet signaling and responses in wild-type (WT) and Syk(Y130F) littermates. Platelets from homozygous Syk(Y130F) mice showed a decrease in functional responses after activation with CRP, a GPVI agonist, and CLEC-2 crosslinking compared to WT littermates with no significant differences in responses to PAR-4 or purinergic receptor agonists. Key signaling events triggered *via* both GPVI and CLEC-2, including phosphorylation LAT and PLC*γ*-2, were also reduced in Syk(Y130F) platelets at low agonist concentrations. Consistent with these findings, the time to occlusion in the FeCl_3_ injury model and bleeding time in the tail bleeding assay were significantly enhanced in Syk(Y130F) mice compared to WT littermates. Thus, phosphorylation of Syk Y130 enhances GPVI- and CLEC-2-mediated signaling and functional responses in platelets affecting thrombosis and hemostasis.

## Introduction

Spleen tyrosine kinase (Syk) is a 72-kDa non-receptor cytoplasmic protein tyrosine kinase, which serves as an essential component of signal transduction machinery coupled to immune receptor complexes (1–4). Besides the high expression in hemopoietic cells, it is also present in other cells and tissues, such as fibroblasts, epithelial cells, breast tissue, hepatocytes, neuronal cells, and vascular endothelial cells (1, 5). Syk is a crucial signaling protein involved in multiple pathways that lead to downstream events characteristic to individual cell types, including proliferation, differentiation, and phagocytosis (6). Therefore, Syk provides an attractive target for therapeutic intervention in cancer as well as autoimmune and inflammation diseases (7–14).

Syk is composed of distinct functional domains. The major structural regions of Syk are the N-terminal region with two tandem Src-homology-2 (SH2) domains separated by the helical domain termed interdomain A followed by the connecting linker termed interdomain B and finally by the C-terminal catalytic kinase domain. This modular design enables Syk to recognize phosphorylated receptors and transduce signals to downstream effectors with remarkable specificity. The tandem SH2 domains provide high-affinity binding to phosphorylated tyrosine residues within immunoreceptor tyrosine-based activation motifs (ITAMs), which are found in the cytoplasmic tails of various immune receptors (15–19). The ITAM sequences, containing two YXX(L/I) motifs, are present, for instance, in the signal-transducing subunits of B-cell receptor (BCR), T-cell receptor (TCR), and glycoprotein VI (GPVI), as well as in the cytoplasmic tail of the FcγRIIA receptor (15, 20). The hemITAMs, containing a single YXX(L/I) motif, are present in C-type lectin-like receptors, such as CLEC-2, Dectin-1 and -2, which have also been shown to signal in a Syk-dependent fashion (21–25).

Signaling functions of Syk are regulated by its multiple tyrosine phosphorylation (pY) sites (26–34). One key regulatory pY-site is located in the activation loop inside the Syk kinase domain (28, 29, 35, 36), but the density of regulatory pY-sites, including pY317, pY342, and pY346, is especially high within interdomain B(31, 34, 37–45), consistent with the importance of this linker region for the transition of Syk from the autoinhibited to activated conformation (27). Likewise, tyrosine 130 (Y130) is localized to interdomain A, the linker between the two SH2 domains of Syk, and has been shown to be phosphorylated both in cells and in *in-vitro* kinase assays (46–51).

Considering that the location of Y130 may enable pY130 to regulate the interactions of Syk with phosphorylated ITAMs (pITAMs), several studies had been conducted using site-directed mutagenesis in which Y130 was replaced with phenylalanine (Y130F) to prevent phosphorylation at this site. The initial study examining the effect of Y130F in the chicken DT40 B-cell line demonstrated that blocking phosphorylation of Syk Y130 increases co-immunoprecipitation of Syk with BCR, an ITAM-bearing receptor, and BCR-induced phosphorylation of cellular proteins, albeit phosphorylation of Syk itself appears to remain constant (49). However, in the rat RBL-2H3 basophilic cell line, Y130F affected neither binding of Syk to the ITAM-bearing receptor FcεRI, phosphorylation of Syk itself and its key substrates, nor FcεRI-induced cellular responses (50). The latter study is consistent with the finding that the Y130F mutation does not modify binding of Syk expressed in mast cells to a pITAM-containing peptide (52). Therefore, the effect of Y130F mutation on Syk signaling functions appears to be dependent strongly on the cellular context.

Considering our interest in platelet biology, we focused on evaluating the role of Syk Y130 in platelets, anucleate cells that are produced by megakaryocytes whose major function is to respond to vascular damage and mediate hemostasis (53–55). Platelets represent an advantageous experimental system for examining the role of Syk Y130 in primary cells, because Syk is essential for platelet signaling and biological responses (56–59). GPVI, an ITAM-bearing collagen receptor, and CLEC-2, a hemITAM-bearing receptor, are major platelet receptors whose functions and modes of signaling initiation are distinct, but both are critical for platelet functions (20–24, 59–64).

To conduct this study we generated, using CRISPR/Cas approach, knock-in (KI) mice carrying the gene encoding Syk containing the Y130F mutation rendering this kinase incapable of phosphorylation on Y130. Using these mice, we investigated ITAM- and hemITAM-dependent signaling and physiological responses in Syk(Y130F)-expressing platelets, which represent a research approach superior to the use of cell lines reconstituted with wild-type (WT) or mutant Syk.

## Material and methods

All reagents were purchased from Thermo Fisher Scientific unless otherwise stated. Chrono-lume used for the detection of secreted ATP was purchased from Chrono-log corporation (Havertown, PA). Anti-pSyk Y525/526 (mouseY519/520; #2710), anti-pPLC γ 2 Y1217(#3871) anti-pSyk Y352 (mouse Y346; #2701), and anti-pLAT Y220 (#3584) were purchased from Cell Signaling Technology (Danvers, MA). Anti-Syk (#sc-51703) and anti-PLCγ2 (#sc-5283) were purchased from Santa Cruz Biotechnology. Anti-LAT (#TA327925) was purchased from Origene (Rockville, MD). Anti-pSyk 130 was from Dr. Coopman’s lab (65). Blocking buffer was purchased from Genesee Scientific and secondary antibodies (IRDye 800CW goat anti-rabbit and IRDye 680LT goat anti-mouse) were purchased from Li-Cor (Lincoln, NE). CRP-A was purchased from Pplus Medical (Salisbury, United Kingdom). The CLEC-2 activating antibody was purchased from Biolegend (San Diego, CA). AYPGKF peptide was purchased from GenScript (Piscataway, NJ).

### Generation and housing of Syk(Y130F) mice

Syk (Y130F) KI mice were generated using the CRSPR/Cas9 technique on the C57BL/6 background by Taconic/Cyagen (Santa Clara, CA). The Syk Y130F mutation was confirmed by both sequencing and digesting the PCR product with the restriction enzyme, EcoRI. The oligonucleotides used for PCR are forward primer 5’-*CTACACCATCGAGAGGGAACTTAAT-3’ and* reverse primer 5’*GTTAAAACACAACCTTCTCCTTCGT-3’*. Mice were housed in a specific pathogen-free facility, and all animal procedures were approved by the Temple University Institutional Animal Care and Use Committee (protocol #4864).

### Platelets preparation

Mouse blood was collected, and platelets were isolated as described previously (66).The isolated platelets were counted using a Hemavet 950FS blood cell analyzer and adjusted to a final concentration of 1.5x10^8^ cells/ml in a modified Tyrode’s solution (138mM NaCl, 2.7mM KCl,2-mM MgCl2, 0.42mM NaH2PO4, 10mM HEPES, 0.1% glucose, and 0.2U/mL apyrase, pH 7.4). All platelet aggregation and secretion experiments were carried out using a lumi-aggregometer (Chrono-log) at 37°C under stirring conditions. Platelet aggregation was measured using light transmission, and ATP secretion was measured using Chrono-lume (a luciferin/luciferase assay).

### Western blotting

Western blotting procedures were performed as described previously (66). Briefly, platelets were stimulated for 3 minutes with CRP or for 5 minutes with the CLEC-2 Ab with stirring at 37°C. The reaction was stopped by precipitating the platelet proteins with HClO_4_ at a final concentration of 0.6 N. The pellet was washed once with deionized water prior to the addition of sample loading buffer. Platelet protein samples were then boiled at 95°C for 5 minutes prior to resolution by SDS-PAGE. Proteins were transferred to nitrocellulose membranes using an iBlot (ThermoFisher) for 10 min at 20v. Membranes were blocked with Prometheus OneBlock Western-FL blocking buffer (Genesee Scientific) and incubated overnight at 4°C with primary antibodies against the indicated protein. The membranes were then washed 4x5 min with Tris-buffered saline containing 0.1% Tween-20 (TBST) prior to incubation with appropriate secondary antibodies for 1 hour at room temperature. The membranes were washed again as above and scanned using a Li-Cor Odyssey infrared imaging system.

### Flow cytometry

Isolated platelets (10^6^/100µl modified Tyrode’s buffer/test) were stained for the surface receptor GPVI using 5µl anti-GPVI (Emfret analytics # M011-1) or isotype control IgG-FITC (Emfret Analytics# P190-1), or CLEC-2 using 0.4 mg anti-CLEC-2-PE (Biolegend #146103) or PE-conjugated rat IgG2_b,_**_κ_** (BD Pharmingen# 556925), and kept at room temperature in the dark for 15 min, then fixed with 1% paraformaldehyde in PBS.

To determine platelet P-selectin exposure or αIIbβ3 activation after stimulation with different concentrations of CRP and CLEC-2 Ab, isolated platelets were diluted in Tyrode’s buffer containing 1-2mM CaCl_2_ (10^6^/test), then stimulated with the agonist for 15 min at room temperature. The platelets were then stained with anti-CD41 (BioLegend# 133914), anti-CD62P P-selectin-FITC (Emfret# D200), and anti-JON/A-PE (Emfret# D200) at room temperature for 15 minutes in the dark. The reactions were stopped by addition of 500µl 1% paraformaldehyde in PBS to each tube.

For measuring annexin V binding, isolated platelet samples were diluted in 1x Annexin V binding buffer (BD Pharmingen# 51-66121E) with 2mM MgCl_2_ and 0.2U/mL apyrase, then stimulated for 20 minutes at room temperature with different concentrations of CRP. The isolated platelets (10^6^/100µl buffer) stained with anti-CD41-BUV395 (BD Bioscience# 752966) and AnnexinV-AlexaFluor647 (BD Pharmingen# 567356). The reactions were stopped by addition of 1ml of 1x Annexin V binding buffer and samples were acquired immediately with 50x10^3^ genuine platelets events were recorded based on CD41 expression.

To assess the platelets leukocytes aggregation, 200µL of whole citrated blood was kept to rest for 10 min then stained with anti-CD45-AlexaFluor700 (Invitrogen# 56-0451-80), anti-CD61-PE (BD Bioscience# 561910), anti-Ly6G-BV421(Biolegend# 127627), anti-Ly6C-APC (Invitrogen# 17-5932-82), anti-CD3-BUV737 (Fisher Scientific# 36-700-3280), anti-CD11b-Pe-Cy7 (Invitrogen# 25-0112-82), anti-CD19-BV605 (Invitrogen# 406-0193-80), and anti-P-selectin-FITC (Emfret# D200). At the same time, 20µg/mL of CRP was added and samples kept at RT in the dark for 25 min. Then, 2ml of 1x RBCs lysis/fixation was added per tube for 20 min at RT in the dark. Samples were acquired immediately at low flow rate with recoding of at least 30x10^3^ CD45^+^CD61^+^ events.

FACSymphony A5 flow analyzer was used for data acquisition, and FlowJo was used for data analysis.

### Tail-bleeding assay

Tail bleeding times were determined according to the procedure described by Patel et al (67). Mice aged 4 to 6 weeks were anesthetized prior to amputation of the distal 3 mm of the tail. The tail was then immersed in 37°C saline, and bleeding was monitored. The bleeding was halted manually by applying pressure if it continued longer than 600 seconds.

### Immunoprecipitation

Platelets from WT and Syk Y130F mice were isolated as described above and the count was adjusted to 5 x 10^8^/ml. Aliquots of 225 ul were equilibrated at 37℃ in an aggregometer with stirring and either or left unstimulated or activated with CRP-A. Reactions were stopped with 25 µl 10x TNE lysis buffer (1x: 10 mM TRIS, 150 mM NaCL, 10 mM EDTA, pH 7.4 containing 1% NP-40 and protease/phosphatase inhibitors). Samples were rocked at 4℃ for 30 minutes and centrifuged at 12,000g/4℃/10 minutes. Supernatants were transferred to clean tubes and 5 µg anti-Syk (D3Z1E; Cell Signaling) added. Samples were rocked 60 minutes at 4℃ and 50 µl TruBlot anti-rabbit Ig IP agarose beads (Rockland, cat # 00-8800-25) added. Samples were rocked overnight at 4℃. Beads were pelleted, washed 3 times with 1x TNE lysis buffer, and 50 µl 2x sample buffer added. All samples were boiled. Samples were run of SDS-PAGE as described above.

### Carotid artery injury

FeCl_3_ was used to injure the carotid artery as previously described (67). Mice aged 10 to 12 weeks were anesthetized, and the carotid artery was exposed. A baseline blood flow reading was obtained using a Transonic T402 flow meter. The carotid artery was injured using a 1x1 mm piece of filter paper saturated with 7.5% FeCl_3_ for 90 seconds. The filter paper was removed, and blood flow was recorded.

### Statistics

Data are presented as mean ± SEM of at least three independent experiments. Shapiro–Wilk test is used to assess the normality of the data. Means were compared by: Welch’s test for comparison of two groups; one-way ANOVA for comparison of more than two groups; and two-way ANOVA for comparison of different groups with different factors followed by Tukey *post hoc* multiple comparison test using GraphPad Prism, v.10 (GraphPad Software, Inc., San Diego, CA, USA) where *p*<0.05 was considered statistically significant.

## Results

### Generation and characterization of Syk Y130 knock-in mice

To determine the role of Syk Y130 phosphorylation in ITAM and hemITAM receptor-mediated platelet signaling and responses, we generated a Syk (Y130F) knock-in mouse using the CRSPR/Cas9 approach (Figure 1A). The Syk (Y130F) mutation was confirmed by both sequencing and digesting the PCR product with the restriction enzyme EcoRI. The Syk knock in mutant contains a site for EcoRI cleavage but not the WT allele. After digestion, the following products were detected: WT (+/+) 439 bp, heterozygous (+/−) 239, 200 and 439bp, and homozygous KI (−/−) 239, 200 bp (Figure 1B). In addition, Sanger sequencing of PCR products of WT and Y130F homozygous mouse DNA shows that the relevant codon has been mutated from TAT to TTC, changing a tyrosine to a phenylalanine (Figure 1C). To confirm the Y130F mutation at the protein level, we isolated platelets from Syk(Y130F) and WT littermate mice and stimulated them with different concentrations of collagen-related peptide (CRP), then Syk was immunoprecipitated. The phosphorylation of Y130 was evaluated in the immunoprecipitates using Western blot analysis. As shown in Figure 1D, there was no phosphorylation of Y130 in CRP-treated Syk(Y130F) platelets, while WT samples exhibit a robust response. Together, these data demonstrated that we successfully generated Syk (Y130F) KI mice. Both heterozygous and homozygous Syk Y130F mice bred normally and produced pups at expected Mendelian ratios. Blood cell counts were not significantly altered in Syk (Y130F) KI mice (Table 1). This represents the first *in-vivo* model harboring a mutation at the Y130 site of Syk.

**Figure 1.**
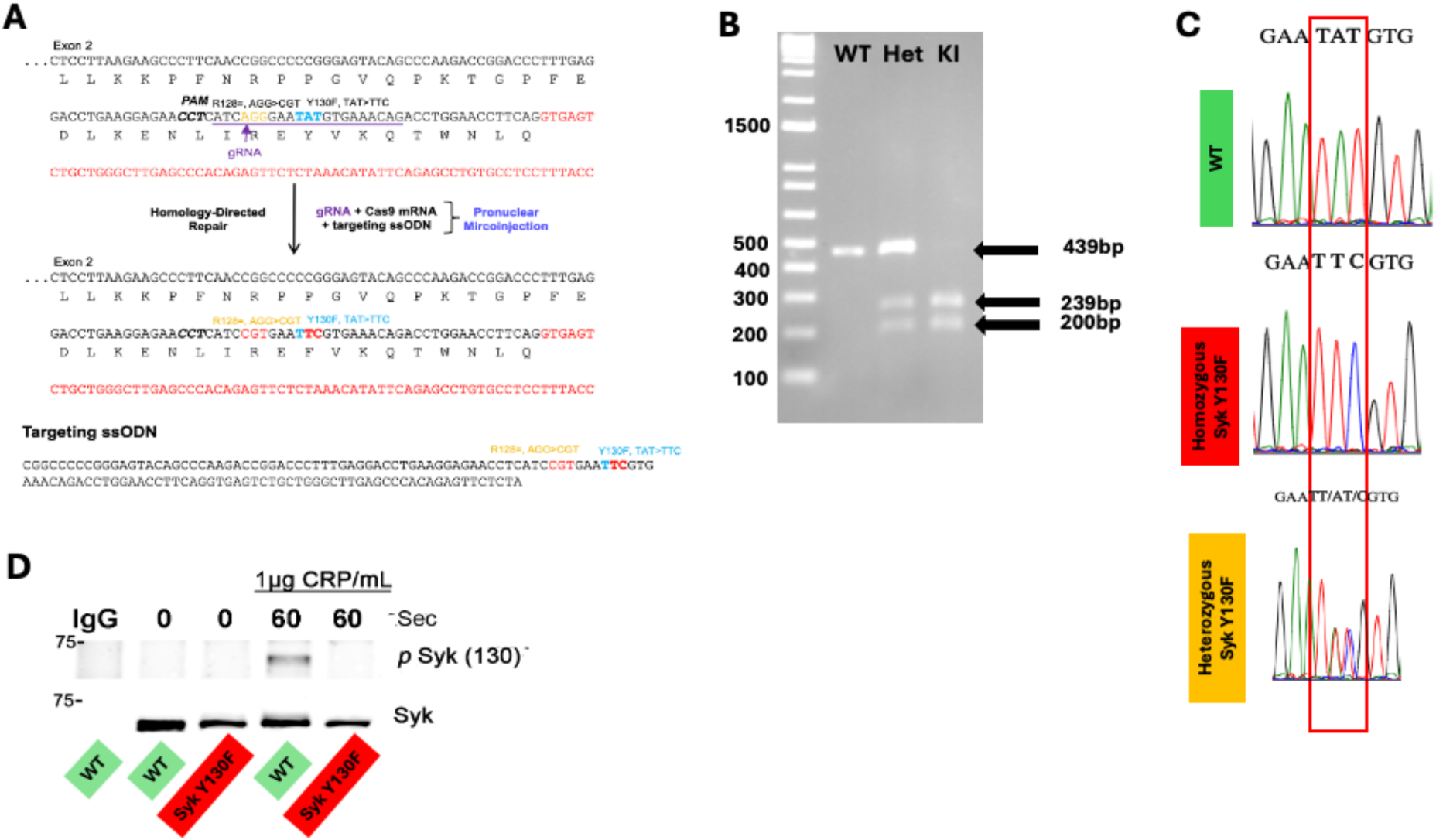
Generation of Syk Y130 knock-in mice. A) the process of creating the mutant mouse via CRISPR is schematically presented. The targeted DNA codon is changed from TAT to TTC, changing the amino acid from tyrosine (Y) to phenylalanine (F). B) Restriction enzyme *Eco*RI digest of PCR products from Syk(Y130F) homozygous knock-in ^(SykY130F/Y130F)^, Syk(Y130F) heterozygous ^(SykY130F/WT)^ (Het), and WT littermate control ^(SykWT/WT)^ DNA. The oligonucleotides used for PCR are forward primer 5’-*CTACACCATCGAGAGGGAACTTAAT-3’*and reverse primer 5’*GTTAAAACACAACCTTCTCCTTCGT-3’.* WT PCR product is 439bp while that of knock-in is 239 and 200bp. C) Sanger sequencing of PCR products of WT and Syk(Y130F) homozygous mouse DNA shows that a DNA codon changed from TAT to TTC changing the tyrosine amino acid to phenylalanine. D) Representative Western blot reveals absence of phosphorylated Syk Y130 in Syk(Y130F) mice compared to WT mice after stimulation of platelets with 1µg CRP (collagen-related peptide), followed by immunoprecipitation of Syk, and probing with anti-Syk pY130.

**Table 1.** Blood cell counts from Syk Y130F and WT littermate control mice.

| Sample | WBC(K/mL) | NE (K/mL) | LY<br>(K/mL) | RBC (M/mL) | PLT (K/mL) | MPV (fL) |
| --- | --- | --- | --- | --- | --- | --- |
| WT | 4.75± 1.13 | 0.9 ± 0.31 | 5.81 ± 1.21 | 6.94 ± 0.2 | 688.71 ± 77.11 | 4.5 ± 0.05 |
| SYK Y130F | 5.3 ± 1.19 | 1.09 ± 0.21 | 6.11 ± 0.9 | 7.11 ± 0.3 | 691.50 ± 84.34 | 4.5 ± 0.09 |

Abbreviations: LY, lymphocyte; MPV, mean platelet volume; NE, neutrophil; PLT, platelet; RBC, red blood cell; WBC, white blood cell.

### GPCR-mediated platelet reactivity is intact in Syk (Y130F) mice

We evaluated the platelet function in Syk(Y130F) mice after stimulation of the G protein–coupled receptor PAR-4 with AYPGKF or the purinergic receptors P_2_Y_1_ and P_2_Y_12_ with 2-MeSADP. Figure 2 shows that Syk (Y130F) platelets respond normally to these agonists. Agonist-induced aggregation or secretion were not significantly different in platelets from the KI and WT littermates.

**Figure 2.**
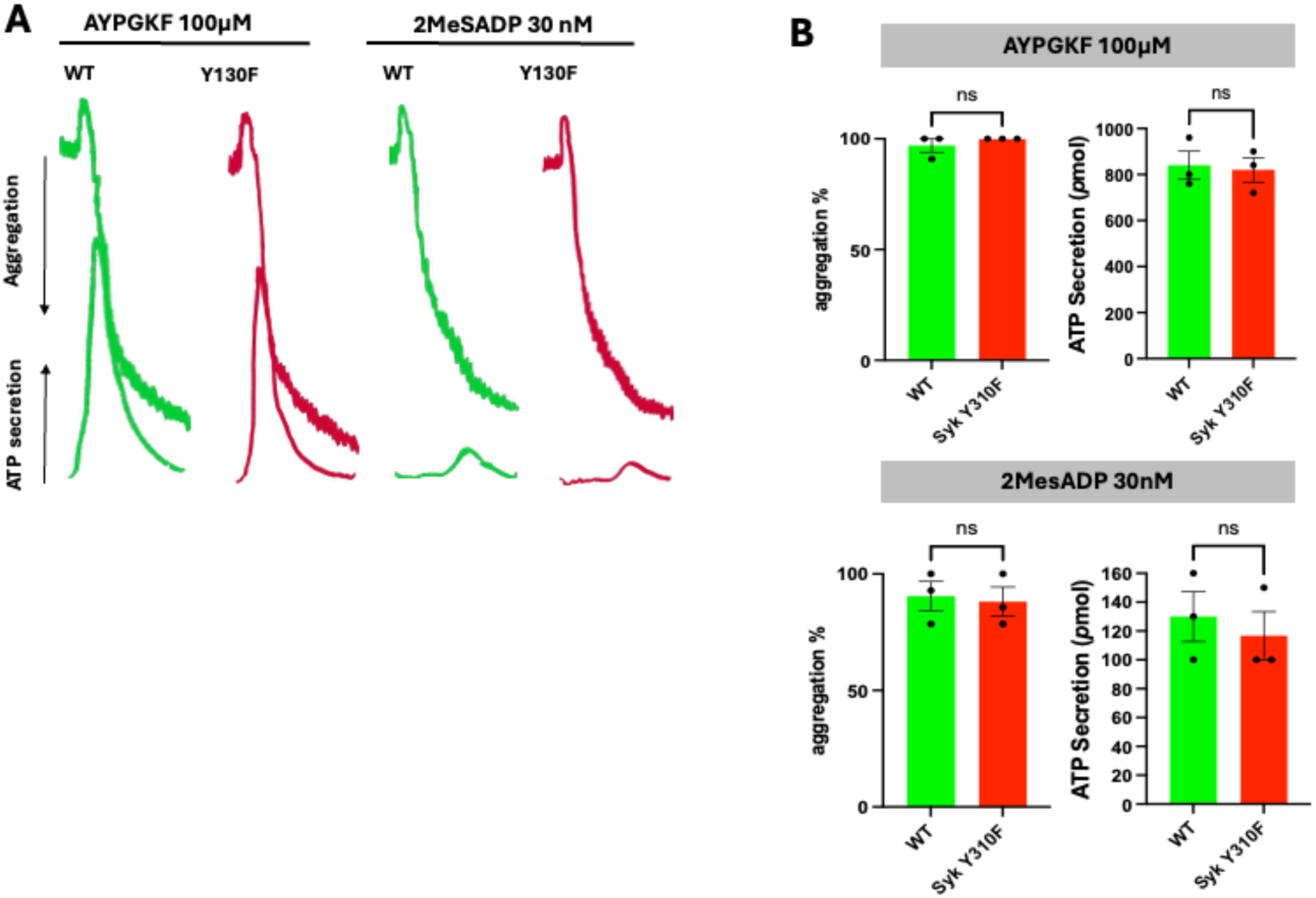
GPCR-mediated platelet reactivity is intact in Syk (Y130F) mice. A) Aggregation and secretion representative tracings of platelets from WT and Syk(Y130F) mice stimulated with the PAR4 agonist AYPGKF (left panel), or P2Y receptor agonist 2-MeSADP (right panel). B) quantitation of aggregation and secretion of platelets from Syk(Y130F) and WT mice stimulated with 100 μM AYPGKF show no significant differences between the two groups. C) quantitation of aggregation and secretion of platelets from Syk(Y130F) and WT mice stimulated with 30 nM 2-MeSADP. The results are representative of at least 3 independent experiments. GPCR, G protein–coupled receptor; Syk, spleen tyrosine kinase.

### Reduced GPVI-mediated aggregation and secretion in Syk(Y130F) platelets

To evaluate the effect of Syk Y130F on GPVI-mediated aggregation and secretion, we stimulated isolated platelets from Syk(Y130F) and WT littermate mice with different concentrations of CRP. Our results indicated that there was no aggregation or ATP secretion in Syk(Y130F) platelets at very low concentrations of CRP (0.125 µg/mL), while WT platelets aggregated and secreted under this condition (Figure 3A and B). Furthermore, Syk(Y130F) platelets showed a significant decrease in ATP secretion in a concentration-dependent manner compared to platelets from WT littermates (Figure 3A and B). As these differences could occur because of altered expression of GPVI on platelets in Syk(Y130F) platelets, we evaluated the GPVI expression on WT and Syk(Y130F) isolated platelets using flow cytometry. The result showed no significant difference between the two groups (Figure 3D). We also evaluated CRP-induced P-selectin exposure, which indicates alpha granule release, and JON/A antibody binding to ⍺IIbβ3, which indicates activation of this integrin. The results shown in Figure 3C indicate that both P-selectin and JON/A levels were significantly reduced in Syk(Y130F) platelets compared to WT control platelets after CRP stimulation in a concentration-dependent manner.

**Figure 3.**
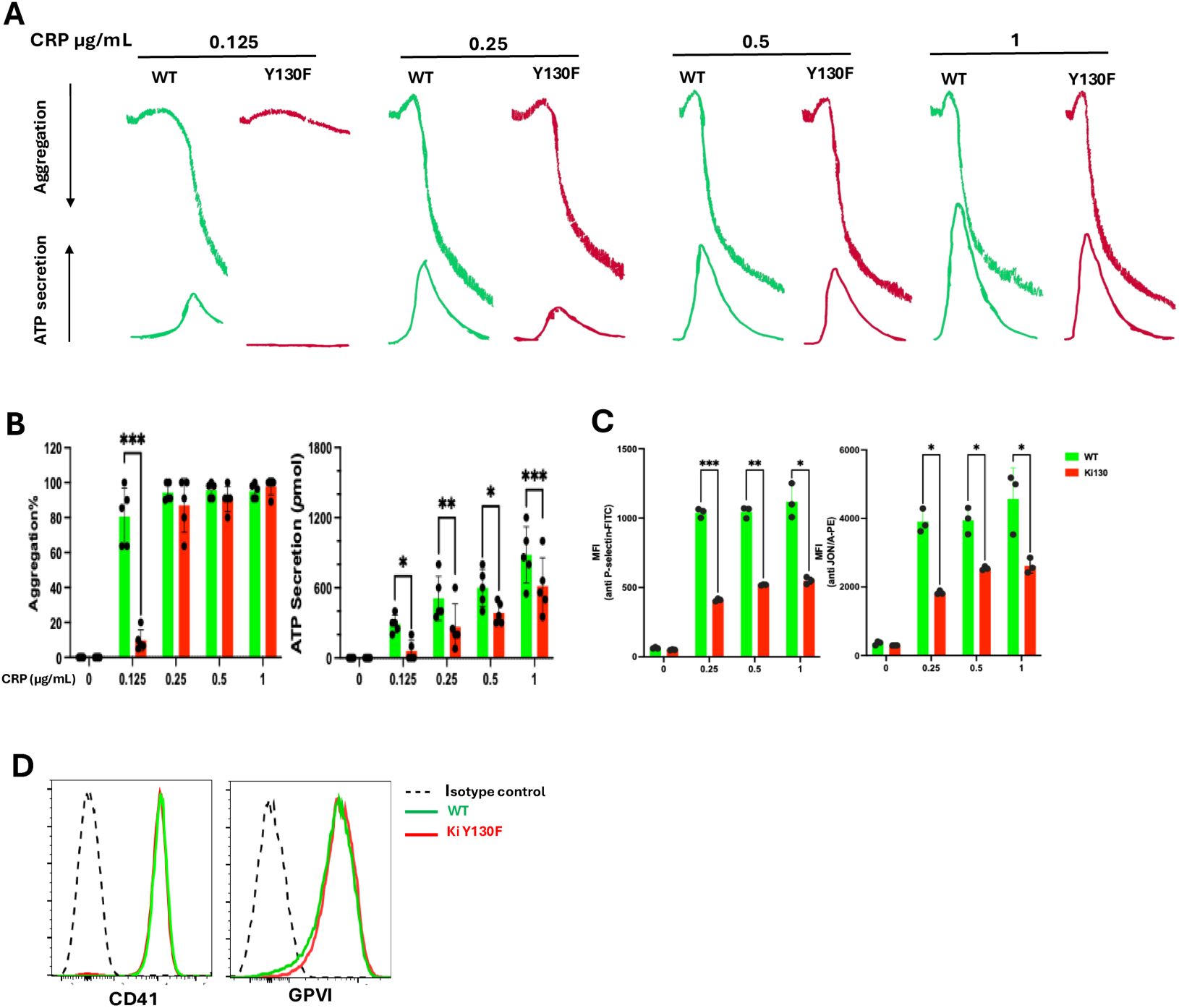
Reduced GPVI-mediated aggregation and secretion in Syk (Y130F) platelets. A) Representative aggregation and secretion tracings of Syk (Y130F) and WT control platelets after using of indicated concentration of CRP as an agonist. B) Aggregation quantification (left panel, and ATP secretion (right panel), and C) quantification of P-selectin (left panel) and JON/A levels (right panel). D) Histograms of fluorescence were determined for isolated platelets. FITC anti-GPVI and APC anti CD41 labeling in Syk(Y130F), and WT were compared with a control FITC-IgG1 and APC-IgG1mouse antibodies, respectively. Results are depicted as mean +/-SEM, and differences are analyzed using Two-Way ANOVA with Tukey *post-hoc* test. *P* values; ∗*P* < 0.05; ∗∗*P* < 0.01; and ∗∗∗*P* < 0.001. The results are representative of 3 independent experiments at least. CRP, collagen-related peptide; GPVI, glycoprotein VI; Syk, spleen tyrosine kinase.

### CRP-induced signaling is diminished in Syk(Y130F) platelets

To evaluate GPVI signaling through the Syk-dependent pathway, we analyzed phosphorylation of the main proteins involved in the GPVI signaling cascade. Our results show less phosphorylation of Syk Y346, Syk Y519/520, LAT Y191, and PLC*γ*2 Y1217 in Syk(Y130F) platelets compared to WT platelets (Figure 4). These data indicate that signaling downstream of GPVI is inhibited in Syk(Y130F) platelets correlating well with reduced GPVI-induced aggregation and secretion.

**Figure 4.**
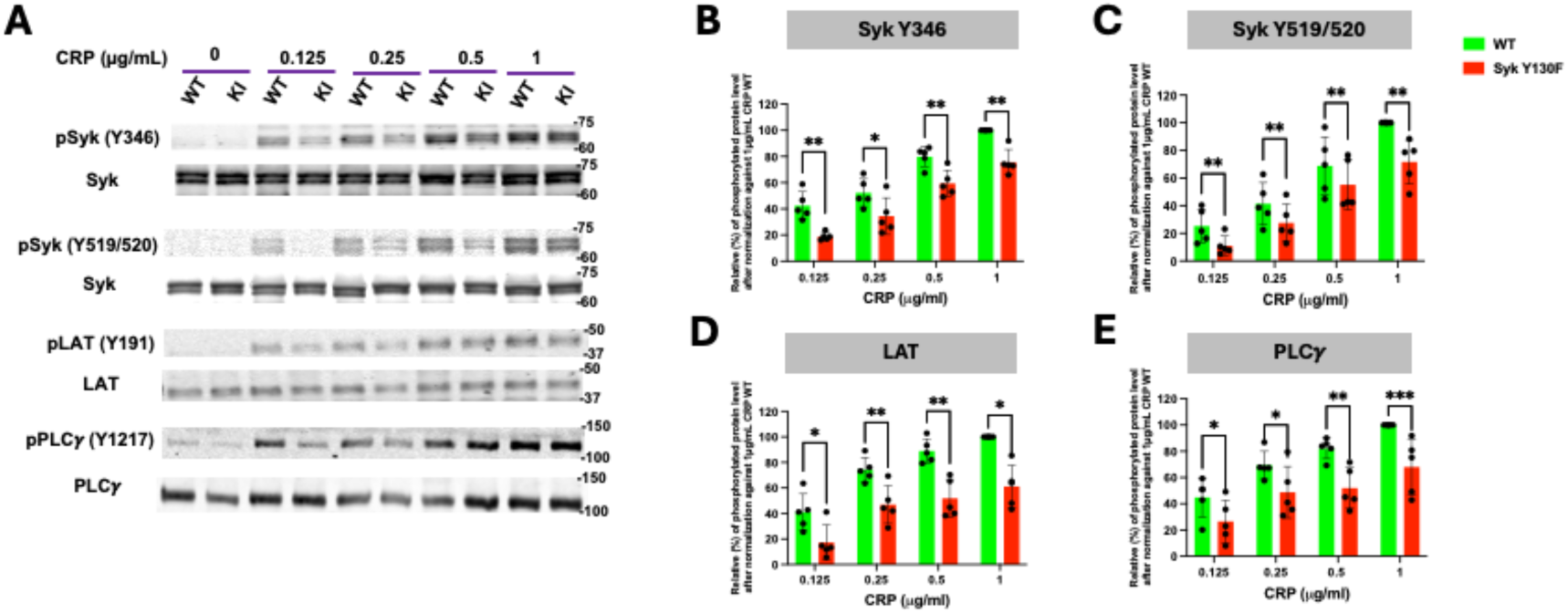
CRP-mediated signaling is diminished in Syk (Y130F) platelets. A) representative Western blots showing the indicated phosphorylated and total protein in Syk(Y130F) and WT littermate control platelets stimulated with the indicated concentrations of CRP for 4 min. **B–E**) quantification of the indicated phosphorylated residue depicted as percentages of WT stimulated with 1µg CRP/mL after normalizing each experiment individually. Results are depicted as mean +/-SEM and differences are analyzed using Two-Way ANOVA with Tukey *post-hoc* test after passing the Shapiro-Wilk normality test. P values; ∗*P* < 0.05; ∗∗*P* < 0.01; and ∗∗∗*P* < 0.001.The results are representative of 5 independent experiments. CRP, collagen-related peptides; Syk, spleen tyrosine kinase.

### Impaired CLEC-2-mediated aggregation and secretion in Syk (Y130F) platelets

We also investigated a role of Syk Y130 phosphorylation in hemITAM-dependent signaling that is initiated through the CLEC-2 receptor by its cross-linking with different concentrations of an anti-CLEC-2 monoclonal antibody. Reduced responses, including aggregation, α-granule secretion, dense granule secretion, and integrin activation, were seen in the Syk(Y130F) platelets as compared to the WT platelets (Figure 5). These results align well with our findings related to GPVI-stimulated platelets responses shown above. As these differences could occur because of altered expression of CLEC-2 on platelets in Syk(Y130F) platelets, we evaluated the CLEC-2 expression on WT and Syk(Y130F) isolated platelets using flow cytometry. The result showed no significant difference between the two groups (Figure 5D).

**Figure 5.**
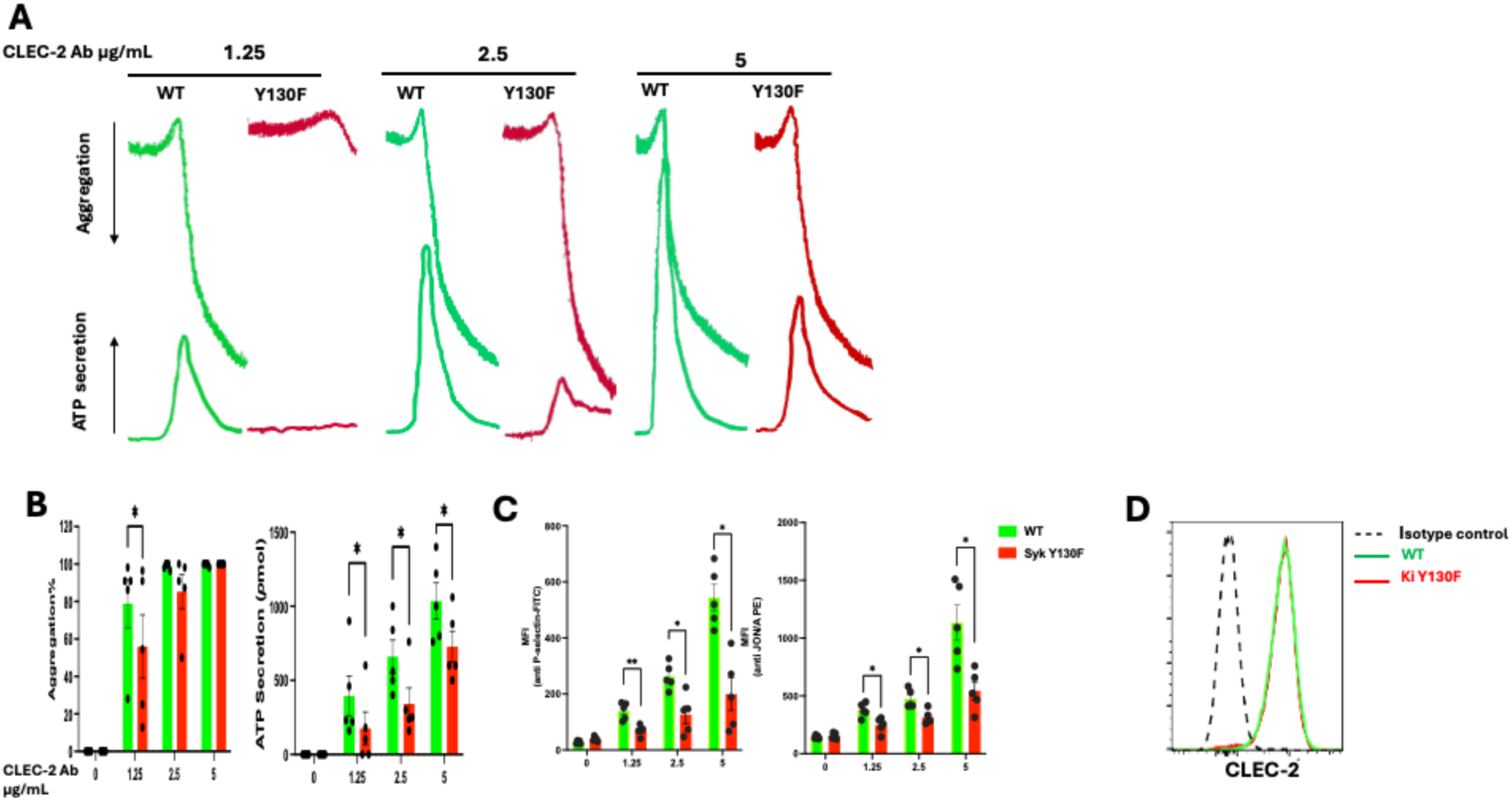
Inhibited CLEC-2-mediated aggregation and secretion in Syk (Y130F) platelets. A) Representative aggregation and secretion tracings of Syk(Y130F) and WT control platelets after using indicated concentration of CLEC-2 Ab as an agonist. B) Aggregation quantification (left panel, and ATP secretion (right panel). C) Surface P-selectin (left panel) and JON/A (right panel) after CLEC-2 Ab stimulation from 5 independent experiments. D) Histogram of fluorescence was determined for CD41^+^ events, PE anti-CLEC-2 labeling in Syk(Y130F), and WT were compared with a control PE-IgG2 rat anti mouse antibody. Results are depicted as mean +/-SEM, and differences are analyzed using Two-Way ANOVA with Tukey *post-hoc* test. P values; ∗*P* < 0.05; and ∗∗*P* < 0.01. The results are representative of 5 independent experiments. CLEC-2, C-type lectin-like type II transmembrane receptor; Syk, spleen tyrosine kinase.

### CLEC-2 -mediated signaling is diminished in Syk(Y130F) platelets

Similar to evaluating Syk signaling cascade downstream of GPVI, an ITAM-bearing receptor, we analyzed phosphorylation of the main signaling proteins involved in the pathway downstream of CLEC-2. Phosphorylation of Syk Y346, Syk Y519/520, LAT Y191, and PLC*γ*2 Y1217 induced through the CLEC-2 receptor was significantly reduced in Syk(Y130) platelets compared to WT platelets (Figure 6).

**Figure 6.**
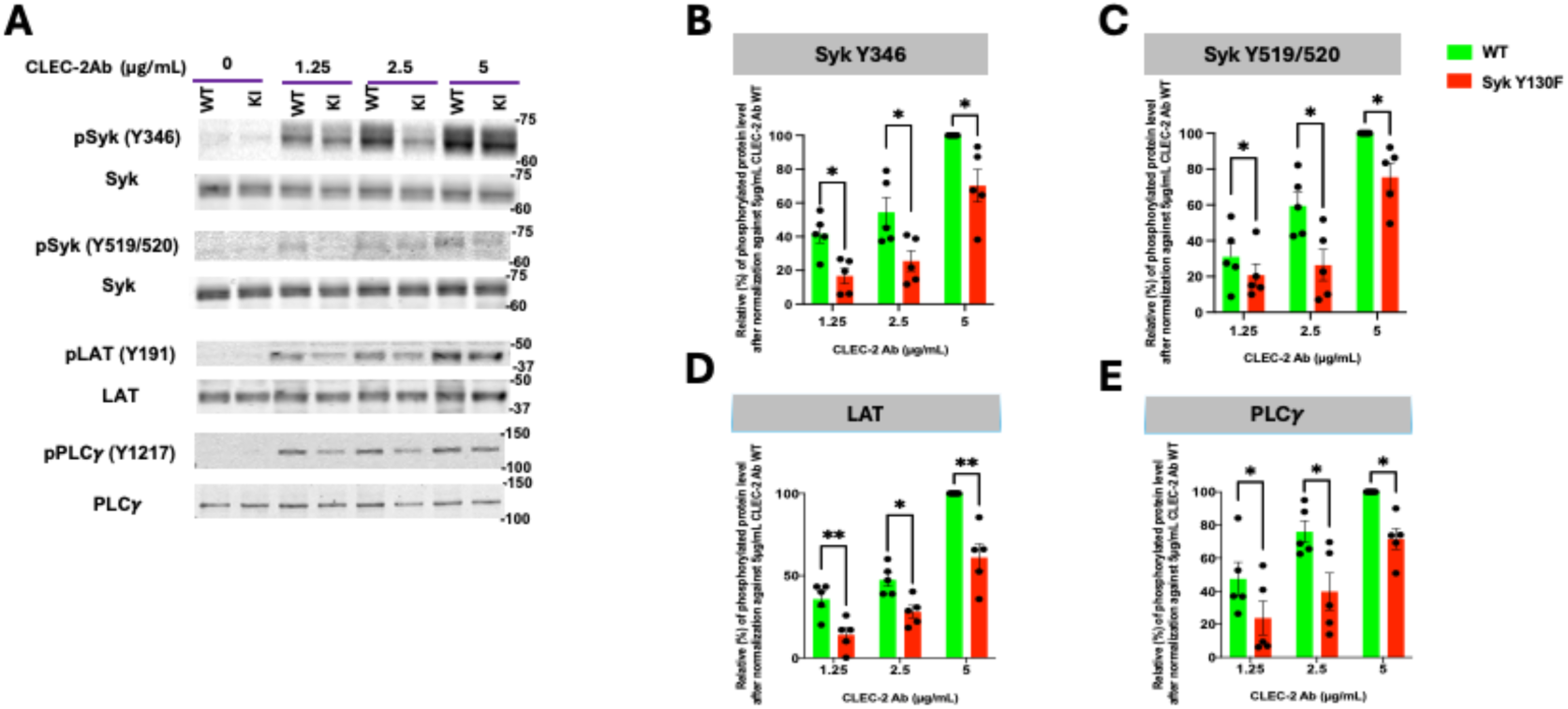
CLEC-2 -mediated signaling is inhibited in Syk (Y130F) platelets. A) Representative Western blots showing the indicated phosphorylated and total protein in Syk(Y130F) and WT littermate control platelets stimulated with the indicated concentrations of CLEC-2 Ab B–E) quantification of the indicated phosphorylated residue depicted as percentages of WT stimulated with 5µg CLEC-2 Ab/mL after normalizing each experiment individually. Results are depicted as mean +/-SE and differences are analyzed using Two-Way ANOVA with Tukey *post hoc* test after passing the Shapiro-Wilk normality test. *P* values; ∗*P* < 0.05; and ∗∗*P* < 0.01. The results are representative of 5 independent experiments. CLEC-2, C-type lectin-like type II transmembrane receptor*; Syk,* spleen tyrosine kinase.

### In vivo thrombus formation and bleeding time are prolonged in Syk(Y130F) mice

To address the impact of Syk Y130F on hemostasis, we performed tail-bleeding assay on litter-matched WT and Syk Y130F) mice. Our data showed a significant increase in the cessation of bleeding in Syk(Y130F) compared to WT mice (Figure 7A). To determine the impact of impaired phosphorylation of Syk Y130 on the thrombus formation, we utilized a FeCl_3_-injury thrombosis model, monitoring the time that is needed for occlusion of the injured carotid artery in mutant and WT mice. Syk(Y130F) mice formed stable occlusion in a statistically significant longer time than WT mice did (Figure 7B).

**Figure 7.**
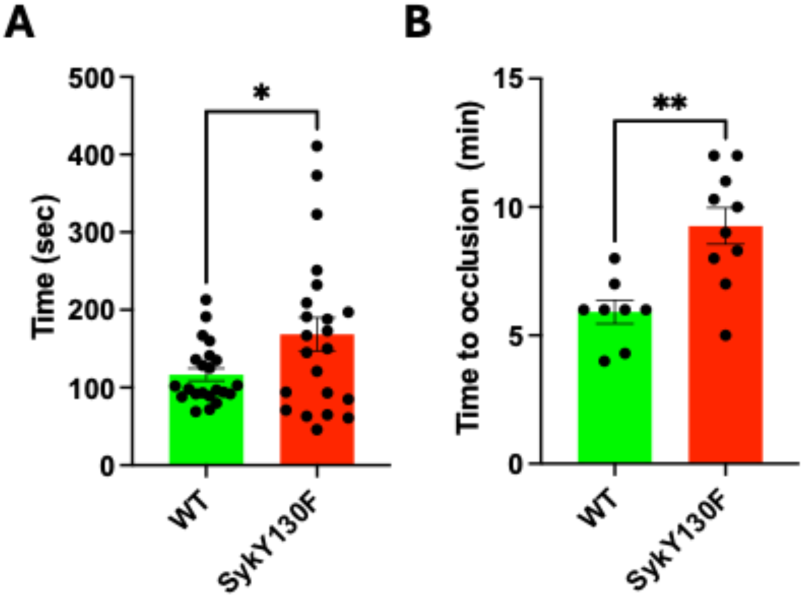
In vivo thrombus formation is prolonged in Syk (Y130F) knock-in mice as well as bleeding time. A) scatter dots plot shows the time it took for bleeding to stop during tail bleeding experiments conducted on homozygous Syk(Y130F) KI and WT littermate control mice in a blind fashion. B) scatter dots plot of the time to occlusion in WT and Syk(Y130F) KI mice following 7.5% FeCl_3_ injury on the carotid artery in a blind fashion. Welch’s test was used to analyze the differences between the two groups after passing the Shapiro-Wilks normality test. *P* values; ∗*P* < 0.05; and ∗∗*P* < 0.01. Syk, Spleen tyrosine kinase.

### Reduced annexin V binding and platelet-leukocytes aggregation in Syk(Y130F) mice compared to WT

Measuring annexin V binding as a marker of phosphatidylserine (PS) exposure which is representative of platelet procoagulant activity, we showed that no differences between resting platelets in both groups. However, after stimulation with different doses of CRP both intensity of florescence with Annexin V and percentages of Annexin V positive platelets (Figure 8A) decreased in Syk Y130F) compared to WT mice in a concentration-dependent manner.

**Figure 8.**
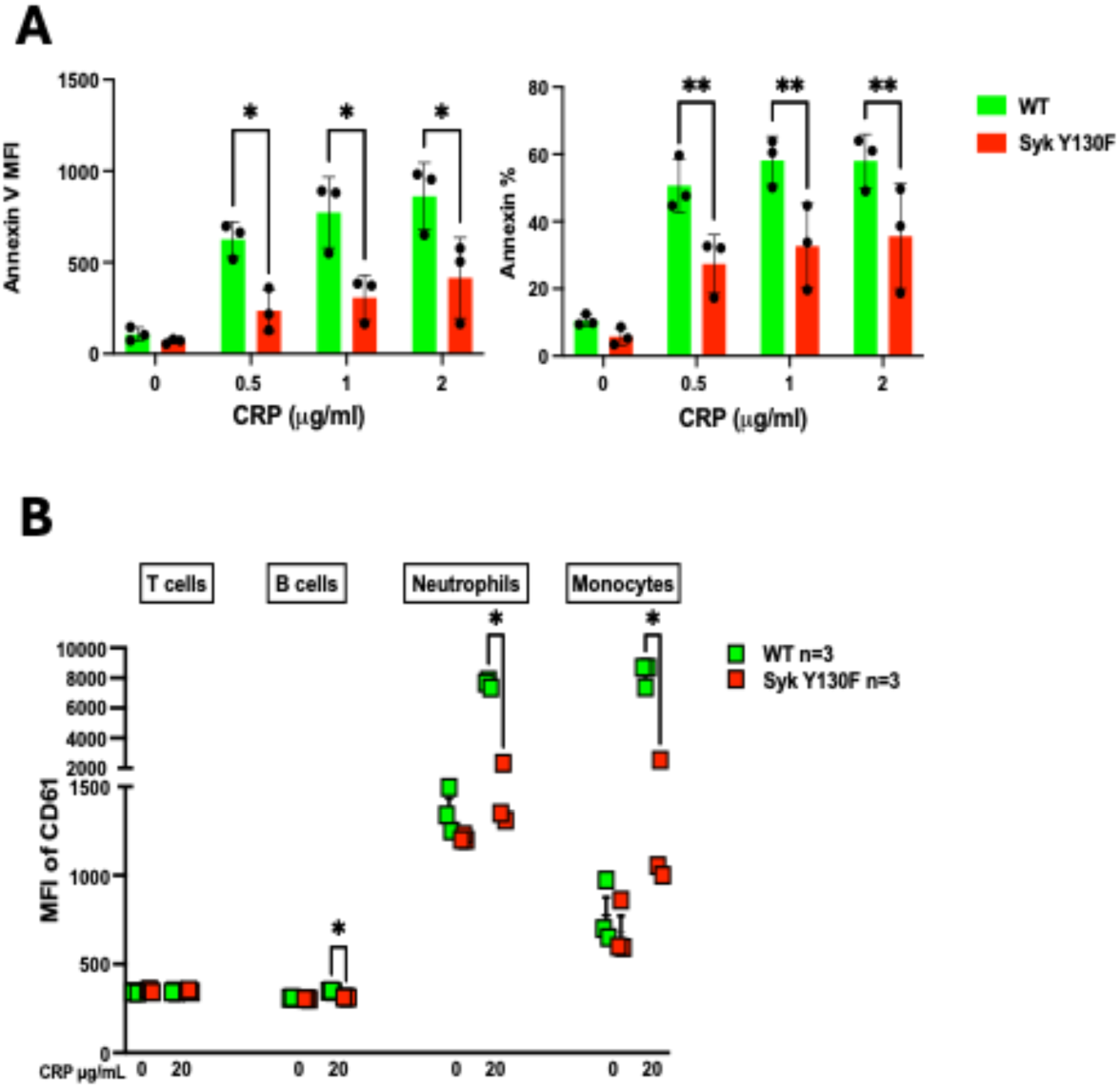
Diminished Annexin V binding and platelets leukocytes aggregation in Syk Y130F mice compared to WT. A) Bar graphs represent mean fluorescence intensity (MFI) and percentage of Annexin V+ platelets with different doses of CRP stimulation, respectively. Results are depicted as mean +/-SEM, and differences are analyzed using Two-Way ANOVA with Tukey *post-hoc* test. B) Scattered dot plots of MFI of CD61 intensities in T cells, B cells, neutrophils, and monocytes indicating platelets leukocytes aggregations with or without 20µg CRP/mL. *P*-values were determined using unpaired Student’s *t*-test. *P*-values are depicted as <0.05 (*). MFI, mean fluorescence intensity.

We also assessed the formation of platelet-leukocyte aggregates (PLAs) across distinct myeloid and lymphoid lineages after stimulation with CRP. There was a marked reduction in platelet aggregation with myeloid cells (monocytes and neutrophils); however, this effect did not extend to the lymphoid compartment, as aggregate levels for both B and T cells remained statistically unchanged (Figure 8B). These data indicate the diminished prothrombotic and proinflammatory status of platelets in Syk(Y130F) mice.

## Discussion

Our previous studies indicated that the signaling functions of Syk in platelets are regulated by its phosphorylation, mostly on tyrosine residues(36, 42, 43, 45, 68). In this study, application of the Y-to-F mutational approach allowed us to examine the role of pY130 in the regulation of Syk in platelets, which has never been evaluated before. Our experiments focused on molecular signaling and biological responses induced through GPVI, a key collagen receptor of platelets, indicated that phosphorylation of Syk pY130 enhances the function of Syk. Indeed, the Y130F mutation blocking phosphorylation of Y130, reduces phosphorylation of (i) Syk on Y519/520 and Y346, which is mostly due, respectively, to autophosphorylation and phosphorylation by Src-family kinases (SFK) (36, 43, 69), (ii) LAT, a critical adaptor protein and a substrate of Syk (70–72), and (iii) PLC-γ2, a key downstream element of the Syk-dependent signaling pathway (57, 71, 73) (Figures 4, 6). Consistent with these effects, Y130F reduces critical platelets responses: secretion, aggregation, α-granule release as measured by the surface exposure of P-selectin, and the ⍺IIbβ3 integrin activation as measured by the antibody recognition of its activated conformation (Figures 3, 5).

The role of pY130 appears to be specific for Syk-dependent processes, since the GPCR-mediated responses have not been modified by Y130F (Figure 2). As expected, this result is consistent with the finding that all Syk Y-to-F mutations we have studied previously modify only Syk-dependent processes in platelets that are induced through ITAM- and hemITAM-bearing receptors, but not platelet responses mediated by GPCR (42, 43, 45, 68).

Notably, although the effect of Y130F on receptor-induced protein phosphorylation and most biological responses typically does not exceed two-fold, it is seen in the entire range of agonist concentrations (Figures 3-6). This result is characteristic for Y130F, since the effects of other Y-to-F mutations that modify Syk functions are typically reduced and even lost when the stimulation strength increases (42, 43, 68). The only response that demonstrates leveling the inhibitory effect of Y130F at high stimulation intensity is aggregation, which generally exhibits not a linear dependence on agonist, but a threshold response pattern. In addition to primary aggregation response, as long as there is some dense granules release the aggregation reaches saturation quickly.

It is of note that Y130F suppresses phosphorylation of both Syk Y346 and Y519/Y520, although the former is phosphorylated mostly by Src-family kinases (SFK), while the latter is mostly autophosphorylated by Syk (36, 43, 69). Hence, pY130 appears to facilitate both types of phosphorylation in Syk, not only autophosphorylation of Syk Y519/Y520 as it may be anticipated from a negative effect of Y130F on Syk enzymatic activity in *in-vitro* kinase assays in previous reports (48, 49).

The effects of Y130F on critical platelet-driven processes *in-vivo* are entirely consistent with its effects on platelet signaling and responses *in-vitro*. First, phosphorylation on Y130 is needed for Syk function and hence lack of phosphorylation suppresses hemostasis, a major platelet function, as evident from the results of tail bleeding assays as well as mechanistically similar artery occlusion indicative of thrombosis (see Figure 7). Second, Y130F disrupts formation of platelet-leukocyte aggregates that are critical for platelet contribution to inflammation, another critical function of platelets (see Figure 8). The reduction in leukocyte platelet aggregates is consistent with reduced P selectin surface expression (Fig 3) as P selectin plays a key role in this aggregate formation. Hence, the effects of Y130F on platelet responses *in vitro* are sufficiently strong to be manifested *in vivo* as well.

The effect of Y130F on cell signaling and responses differs substantially in different cell types. In a chicken B-cell line Y130F apparently enhanced cell signaling, albeit not phosphorylation Syk itself (49), while in a rat basophil line this mutation exerted no effect on either the ITAM-bearing receptor binding or signaling (50). This discrepancy may be due to the differences between individual cell types regarding receptors that trigger Syk-dependent signal transduction and/or signaling/regulatory proteins interacting with Syk. Given the consistency of our findings between the effects of Y130F on various aspects of platelet responses *in-vitro* as well as between those *in-vitro* and *in-vivo*, it may be concluded that in Syk pY130 unequivocally plays a positive role in the platelet regulation of Syk.

The molecular basis of the effect of pY130 on the function of Syk may be two-fold. First, the Y130F mutation decreases the intrinsic enzymatic activity of Syk as measured an *in-vitro* kinase assay using both recombinant Syk and Syk from the lysates of BCR-stimulated cells (48, 49). Second, phosphorylation of Y130 may affect the interaction of Syk with pITAMs. Several studies in which Syk Y130 have been replaced with a residue of glutamic acid (Y130E) to potentially mimic pY130 by introducing a permanent negative charge indicate that this mutation reduces the tandem SH2 affinity to pITAM (51, 74) or increases an off-rate for the binding of Syk to the ITAM-bearing FcεRI receptor (50). This apparently occurs because this negative charge inside the interdomain A partially decouples the two SH2 domains in the tandem (51).

The effect of Y130E, which is intended to mimic pY130, was found to be suppressive not only to the interaction between Syk and pITAM, but also to the signaling function of Syk. Y130E decreased activation-induced Syk-dependent tyrosine phosphorylation of cellular proteins in both DT40 chicken B cells and RBL-2H3 rat basophils (49, 50). These findings seem to be in contrast not only with the results described in our study, but also with the original observations when Y130E caused an increase in Syk *in-vitro* kinase activity, while decreasing the stimulation-induced tyrosine phosphorylation of cellular proteins (49). Interpreting the results obtained with Syk(Y130E), one cannot exclude that while Y130E consistently exerts a *bona-fide* effect on the tandem SH2 structure, which reduces its binding to pITAM, this mutation does not represent pY130 closely enough. Not only a phosphate group attached to an aromatic phenyl group in a phosphotyrosine residue is structurally quite different from a carboxyl group in a short aliphatic chain of glutamic acid, but the fraction of Syk molecules phosphorylated on Y130 appears to be much lower than that of phosphorylated on Y342, Y346 or Y519/520 (48, 51). Namely, only ∼5% of Y130 becomes phosphorylated *in-vitro* in the mix of purified Syk, Lyn, and pITAM, whereas 20-70% of major regulatory sites of Syk do (48). Thus, the introduction of Y130E changes the entire population of Syk, while only a minor fraction of Syk appears to acquire pY130 as a result of cell stimulation. Therefore, the Y130F mutation is likely to represent a more precise tool for probing the role of Syk pY130.

Finally, our results indicate that Syk pY130 affects Syk-dependent signaling downstream of both GPVI, an ITAM-bearing receptor, and CLEC-2, a hemITAM-bearing one (see Figure 3, 4 vs Figure 5,6). Thus, the effect of this pY-site is largely independent of the type of interactions between the Syk SH2 domains and the ITAM/hemITAM of the membrane receptors, despite the clear difference between ITAM and hemITAM in their physical and functional interactions with the Syk tandem SH2 (18, 19, 74, 75) and despite the possible partial decoupling of the two SH2 domains of Syk as a result of Y130 phosphorylation (51). This observation further supports the idea that phosphorylation of Syk Y130 exerts a direct effect on Syk activity and/or functions regardless of its effect on the receptor interactions of Syk SH2 domains.

Taken together with previous findings our current results argue that the existence of Syk pY130 adds another regulatory site to the system governing signaling functions of this kinase, increasing the overall complexity of this multi-layered regulation ensuring appropriately fine-tuned responses to the stimuli of varying strength. In particular, one might speculate that phosphorylation of Syk at Y130 plays a role of a built-in timer mechanism that limits the duration of physical interaction between Syk and membrane receptors thus transitioning Syk from a receptor-bound state to a free cytosolic state and that this transition enhances the function of Syk in platelets.

## Data Availability

All data concerning this report is available in the manuscript.

## Authors statement

All authors have significantly contributed to the article and have read and approved the manuscript.

